# Quantitative measurements of chromatin modification dynamics during zygotic genome activation

**DOI:** 10.1101/601393

**Authors:** Yuko Sato, Lennart Hilbert, Haruka Oda, Yinan Wan, John M. Heddleston, Teng-Leong Chew, Vasily Zaburdaev, Philipp Keller, Timothee Lionnet, Nadine Vastenhouw, Hiroshi Kimura

**Author notes:** Correspondence to: Hiroshi Kimura, Cell Biology Center, Institute of Innovative Research, Tokyo Institute of Technology, 4259 Nagatsuta-cho, Midori-ku, Yokohama, 226-8503, Japan. Present address: Institute of Toxicology and Genetics, Karlsruhe Institute of Technology; Zoological Institute, Systems Biology/Bioinformatics, Karlsruhe Institute of Technology, Germany (Lennart Hilbert). Present address: Department of Biology, Friedrich-Alexander Universität Erlangen-Nürnberg and Max-Planck-Zentrum für Physik und Medizin, Erlangen, Germany (Vasily Zaburdaev).

## Abstract

Histone posttranslational modifications (PTMs) are key gene expression regulators, but their rapid dynamics during development remain difficult to capture. We describe an approach that images fluorescent antibody fragments to quantify PTM enrichment and active transcription at defined loci during zygotic genome activation in living zebrafish embryos. The technique reveals a drastic increase in histone H3 Lys27 acetylation (H3K27ac) before genome activation both at a specific locus and globally.

## Introduction

We have previously developed FabLEM (Fab-based live endogenous modification labeling), a live-cell imaging system able to visualize transcription and histone modification dynamics using modification-specific antigen-binding fragments (Fabs) (Hayashi-Takanaka et al., 2011). FabLEM enables monitoring both the localization and changes in the global level of histone modifications in living cells and embryos, without affecting the cell cycle or developmental processes (Hayashi-Takanaka et al., 2009; Hayashi-Takanaka et al., 2011). The nuclear concentration of Fabs reflects global modification levels in the nucleus because Fabs transiently bind to the target modification and freely pass through nuclear pores due to their small size (Hayashi-Takanaka et al., 2011). Thanks to the signal amplification provided by artificial tandem gene arrays, FabLEM has also provided live-cell kinetics of glucocorticoid-stimulated transcription activation in cultured cells (Stasevich et al., 2014a). However, no method yet has been able to capture the dynamics of arbitrary histone modifications or the transcription machinery at a defined endogenous locus in live developing embryos. Here we demonstrate that the FabLEM principle can be applied to capture single locus dynamics in the developing zebrafish embryo.

As a proof of principle, we chose a hallmark of epigenetic changes during development, the maternal to zygotic transition. In zebrafish, the zygotic genome is kept silenced for the first several cell cycles, after which transcription is activated (Kane and Kimmel, 1993). How this zygotic genome activation (ZGA) is regulated has been investigated for decades (Schulz and Harrison, 2018). Because of the dynamic nature of post-translational modifications of histone proteins, the modifications are thought to play important roles in regulating transcription activation. ChIP-seq analysis using early zebrafish embryos revealed an enrichment of H3 Lys4 trimethylation (H3K4me3) in promoters of developmentally-regulated genes (Lindeman et al., 2010; Vastenhouw et al., 2010). Occupancy of those genes by H3K4me3 prior to ZGA was suggested to pre-pattern developmental gene expression (Lindeman et al., 2011). A recent report has also shown dynamic changes of H3 Lys27 acetylation (H3K27ac) at promoters and enhancers before and after zebrafish ZGA by ChIP-seq (Zhang et al., 2018). However, the spatiotemporal dynamics of transcription and histone modifications remain unclear because appropriate tools are lacking. Here we demonstrate that FabLEM can be applied to living zebrafish embryos in order to reveal the changes in histone modifications and active transcription during ZGA at defined loci.

## Results and Discussion

### Monitoring ZGA

To monitor transcription activity, we used Fabs specific for the elongating form of RNA polymerase II (RNAP2), which is phosphorylated at Ser2 in its C-terminal domain consisting of 52 repeats of the sequence Tyr-Ser-Pro-Thr-Ser-Pro-Ser (RNAP2 Ser2ph). Fluorescently labeled Fabs specific for histone modifications and RNAP2 Ser2ph were injected into 1-cell stage embryos (Fig. 1a). At the 4-cell stage, embryos were mounted onto a light sheet (SiMView) (Royer et al., 2016) or a confocal microscope (FV1000), and fluorescence images were collected from the 8- or 16-cell stage up to 6 hours post fertilization (hpf), during which ZGA occurs. We first examined whether the embryos injected with Fabs develop normally, as do mouse embryos (Hayashi-Takanaka et al., 2009; Hayashi-Takanaka et al., 2011). Embryos injected with Fabs exhibited normal morphology during development, similarly to the controls without injection or injected with buffer alone (Supplementary Fig. 1), suggesting that Fabs injected under the conditions used in this study do not affect early development. When whole embryos were visualized using a SiMView light sheet microscope, Alexa488-labeled RNAP2 Ser2ph-Fabs distributed throughout the cells at the 8-cell stage, consistent with a lack of transcription (Fig. 1b and Supplementary movie 1). In contrast, Cy5-labeled Fabs specific for histone H3 Lys9 acetylation (H3K9ac), which is a broad euchromatin mark, were enriched in nuclei and mitotic chromosomes throughout the early developmental stages. RNAP2 Ser2ph-Fabs became slightly accumulated in nuclei in few cells at 128-cell stage, and then obviously concentrated in almost all nuclei at the 512- and 1k-cell stages, during which ZGA takes place. When closely looking at the distribution of RNAP2 Ser2ph-Fabs in the latter stages, they were concentrated in two foci (Fig. 1b; inset). These foci were likely to contain alleles of miR-430 gene clusters on chromosome 4 (Chan et al., 2018; Giraldez et al., 2006; Hadzhiev et al., 2019; Heyn et al., 2014; Hilbert et al., 2018). miR-430 family genes that play a role for clearance of maternal RNAs are expressed at the highest level among the early transcribed genes (Hadzhiev et al., 2019; Heyn et al., 2014). To confirm this notion, a Cy3-labeled Morpholino oligonucleotide that targets miR-430 transcripts (Hadzhiev et al., 2019) was injected with Alexa488-labeled RNAP2 Ser2ph-Fab into living embryos (Fig. 1c; Supplementary movie 2). In good agreement with previous reports (Chan et al., 2018; Hadzhiev et al., 2019), RNAP2 Ser2ph foci were co-localized with bright miR-430 Morpholino signals in living embryos, suggesting that the nuclear and local concentration of RNAP2 Ser2ph-Fab reports on transcription activity in living zebrafish embryos, even though the sensitivity may not be as high as the transcript-specific Morpholino, which binds stably to the target RNA compared to the transient binding of Fabs.

**Figure 1.**
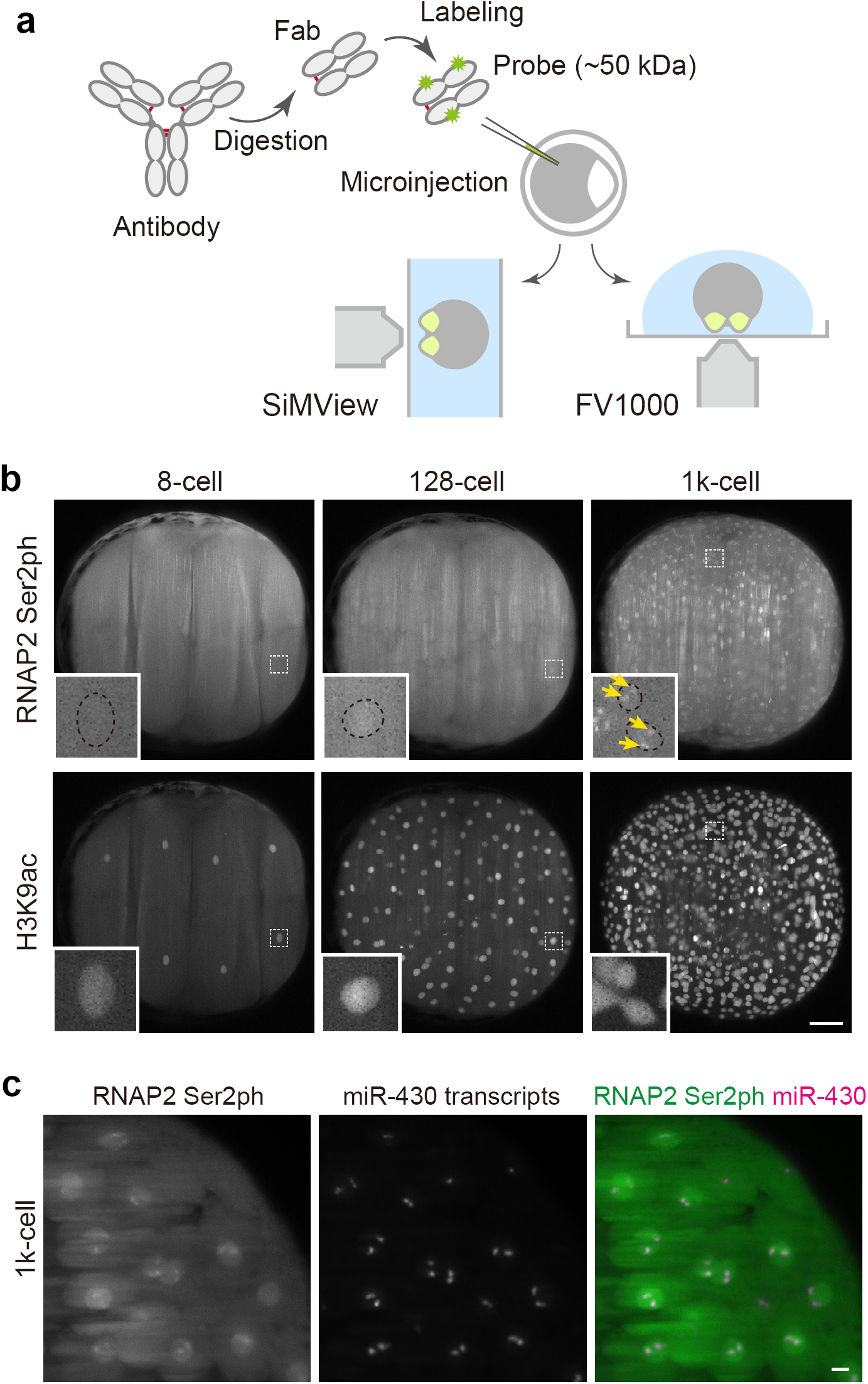
Visualizing RNA polymerase II and histone modifications in living embryo. (a) Scheme of experiments. Fluorescently labeled Fabs prepared from modification-specific antibody are injected into the 1-cell stage zebrafish embryos. After the removal of chorions, the embryos are mounted for a light sheet (SiMView) or a confocal (FV1000) microscope using low-gelling temperature agarose at the 4-cell stage. (b and c) Representative images taken with a SiMView microscope. (b) Fabs specific to RNAP2 Ser2ph (Alexa488) and H3K9ac (Cy5) were simultaneously injected and imaged using SiMView. RNAP2 Ser2ph Fabs were clearly concentrated in nuclei around the 1k-cell stage, whereas H3K9ac Fabs were enriched in nuclei from the 8-cell stage. The maximum intensity projections of 198 z-sections with 2 μm intervals are shown. Insets show magnified views of indicated areas. Yellow arrows in insets indicate RNAP2 Ser2ph foci in nuclei. Scale bar, 100 μm. (c) Co-localization of RNAP2 Ser2ph foci with miR-430 transcripts. Embryos were injected with RNAP2 Ser2ph-Fab (Alexa488) and miR-430 MO (Cy3). Maximum intensity projections (30 z-planes with 2 μm intervals) at 1k-cell stage are shown. Scale bar, 10 μm.

The images acquired with a SiMView feature striping artifacts are common to light sheet microscopes (Power and Huisken, 2017). To quantify how histone modifications change during ZGA, we used a confocal microscope, which yielded a higher spatial resolution and more homogenous background. Time-lapse analysis showed that RNAP2 Ser2ph-Fabs became gradually concentrated in nuclei during embryo development (Fig. 2a), as observed using the light sheet microscope. To quantify the changes in modification levels, we measured the Nucleus/Cytoplasm intensity ratio (N/C ratio) (Hayashi-Takanaka et al., 2011; Stasevich et al., 2014b). The intensity on mitotic chromosomes was often difficult to measure because of their irregular shape and weak Fab enrichments, resulting in poor image segmentation during mitosis (Fig. 2b and Supplementary Fig. 2). We therefore focused on the Fab enrichments in interphase nuclei. The N/C ratio of RNAP2 Ser2ph was slightly increased at the 256-cell stage, and then became higher at the 512- and 1k-cell stages (Fig. 2b; Supplementary Fig. 3), consistently with the timing of zygotic activation (Tadros and Lipshitz, 2009). Again, two bright foci were observed (Fig. 2a; inset at 1k- cell stage). Thus, the nuclear enrichment of RNAP2 Ser2ph-Fabs likely represents the transcription level in zebrafish embryos.

**Figure 2.**
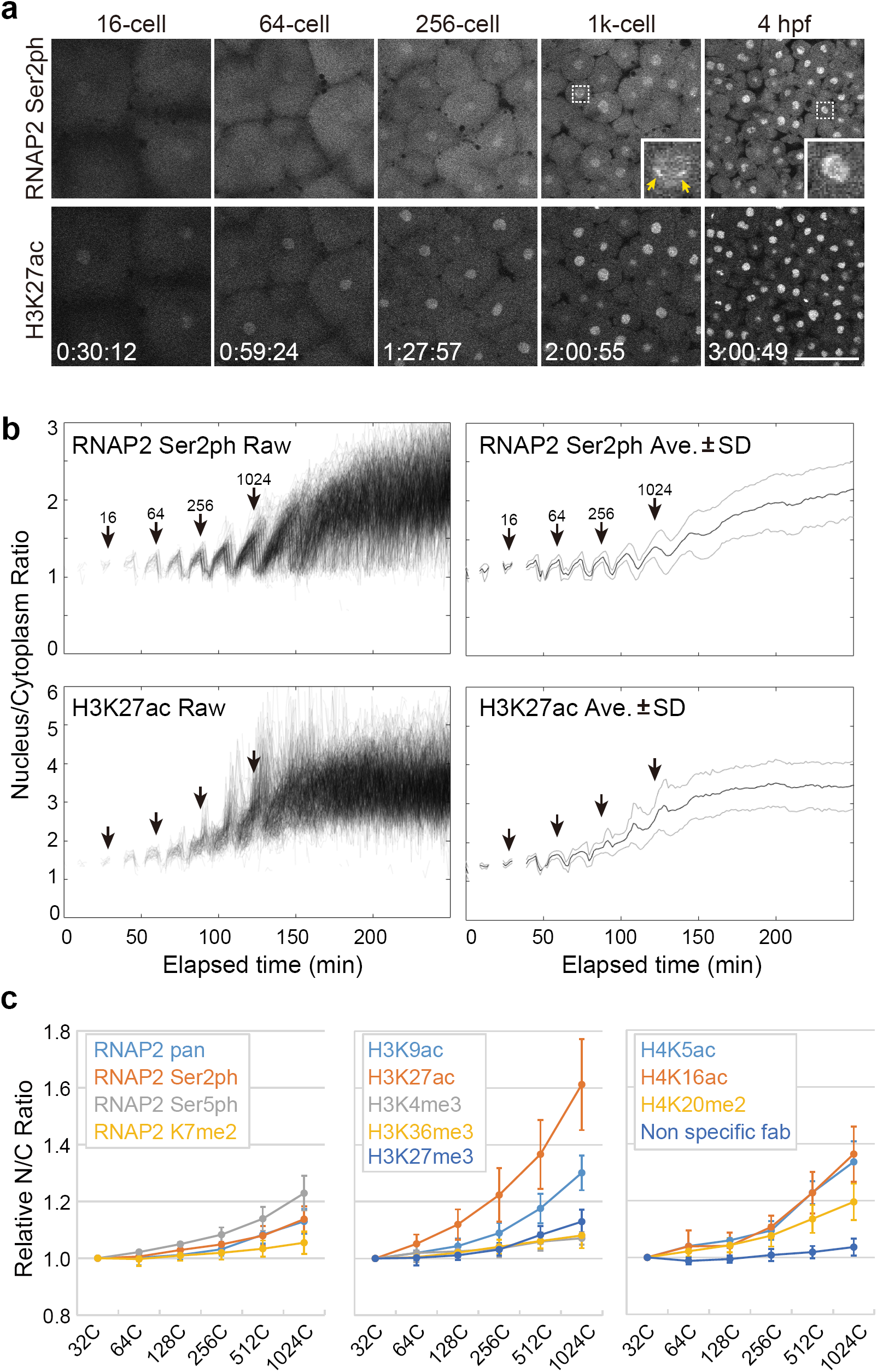
Changes in the level of RNAP2 Ser2ph and histone modifications. (a) Representative images of embryos injected with Fabs specific for RNAP2 Ser2ph (Alexa488) and H3K27ac (Cy3). Insets show magnified views of indicated areas. Yellow arrows in insets indicate RNAP2 Ser2ph foci in nuclei. Elapsed times (h:mm:ss) are indicated. Scale bar, 100 μm. (b) Nucleus/Cytoplasm intensity ratios. Stages judged from time-lapse images are indicated. The global level of H3K27ac in nuclei increased slightly earlier than that of RNAP2 Ser2ph. (c) Relative Nucleus/Cytoplasm (N/C) intensity ratios of RNA polymerase II and histone modifications in interphase cells (averages from 3 independent experiments with standard deviations).

### Quantifying histone modification changes

We next systematically analyzed dynamics of relevant histone modifications. Among the various histone modifications investigated, H3K27ac exhibited the most drastic changes in the early developmental stages. Its nuclear intensity continuously increased from the 64-cell stage, earlier than RNAP2 Ser2ph (Fig. 2a-2c; Supplementary Fig. 3). Acetylation at other sites, like H3K9ac and H4 Lys16 (H4K16ac), and RNAP2 Ser5ph also increased during embryo development, but not as high as H3K27ac during the 64- to 128- cell stages. Subtle increases after the 512-cell stage were also observed for methylation at various sites, including H3 Lys4 trimethylation (H3K4me3), a mark on the transcription start site of actively transcribed gene, H3 Lys27 trimethylation (H3K27me3), a mark associated with facultative heterochromatin, and H3 Lys36 trimethylation (H3K36me3), a mark added with elongating RNAP2 (Fig. 2c; Supplementary Fig. 4). These data are largely consistent with immunofluorescence data using fixed cells (Zhang et al., 2018), and suggest that the dynamic increase of H3K27ac may play a role in transcription activation. H3K4me3 can be a preexisting mark for the target genes (Lindeman et al., 2011; Zhang et al., 2018) but their activation may be mediated through H3K27ac accumulation. Most histone methylations, including H3K4me3, H3K27me3, H3K36me3, and H4K20me2, are gradually established after zygotic transcription (Supplementary Fig. 4), which is probably associated with establishments of epigenomic states depending on cell lineage and differentiation.

### Histone H3K27ac precedes transcription

To clarify the role of H3K27ac on transcription activation, we analyzed the dynamics of H3K27ac during focus formation of RNAP2 Ser2ph during earlier stages (cf. Fig. 2a, inset of the 1k-cell image). To follow the dynamics of RNAP2 Ser2ph and H3K27ac, we acquired images at a higher magnification with shorter time intervals by scanning only a small area just covering a single nucleus from the 64-cell stage. In between the 64- and 128-cell stages, H3K27ac foci appeared in nuclei around the middle of interphase, but the concentration of RNAP2 Ser2ph was hardly observed (Fig. 3b, Supplementary Fig. 5). At the 256-cell stage, H3K27ac foci appeared early in the interphase, and then RNAP2 Ser2ph became concentrated around them (Fig. 3a, b). At 512-cell stage, H3K27ac foci were detected soon after mitosis and RNAP2 Ser2ph foci appeared earlier as well. H3K27ac foci became less intense as RNAP2 Ser2ph foci grew, probably due to chromatin unfolding and/or deacetylation. To confirm the co-localization of H3K27ac and RNAP2 Ser2ph, embryos injected with RNAP2 Ser2ph-Fab were fixed at the 512-cell stage and stained with anti-H3K27ac antibody (Supplementary Fig. 6). As seen in living cells, H3K27ac signals were closely associated with RNAP2 Ser2ph foci. These data indicated that H3K27ac becomes concentrated at miR-430 loci before transcription occurs, suggesting that this histone modification may stimulate transcription in zebrafish embryos.

**Figure 3.**
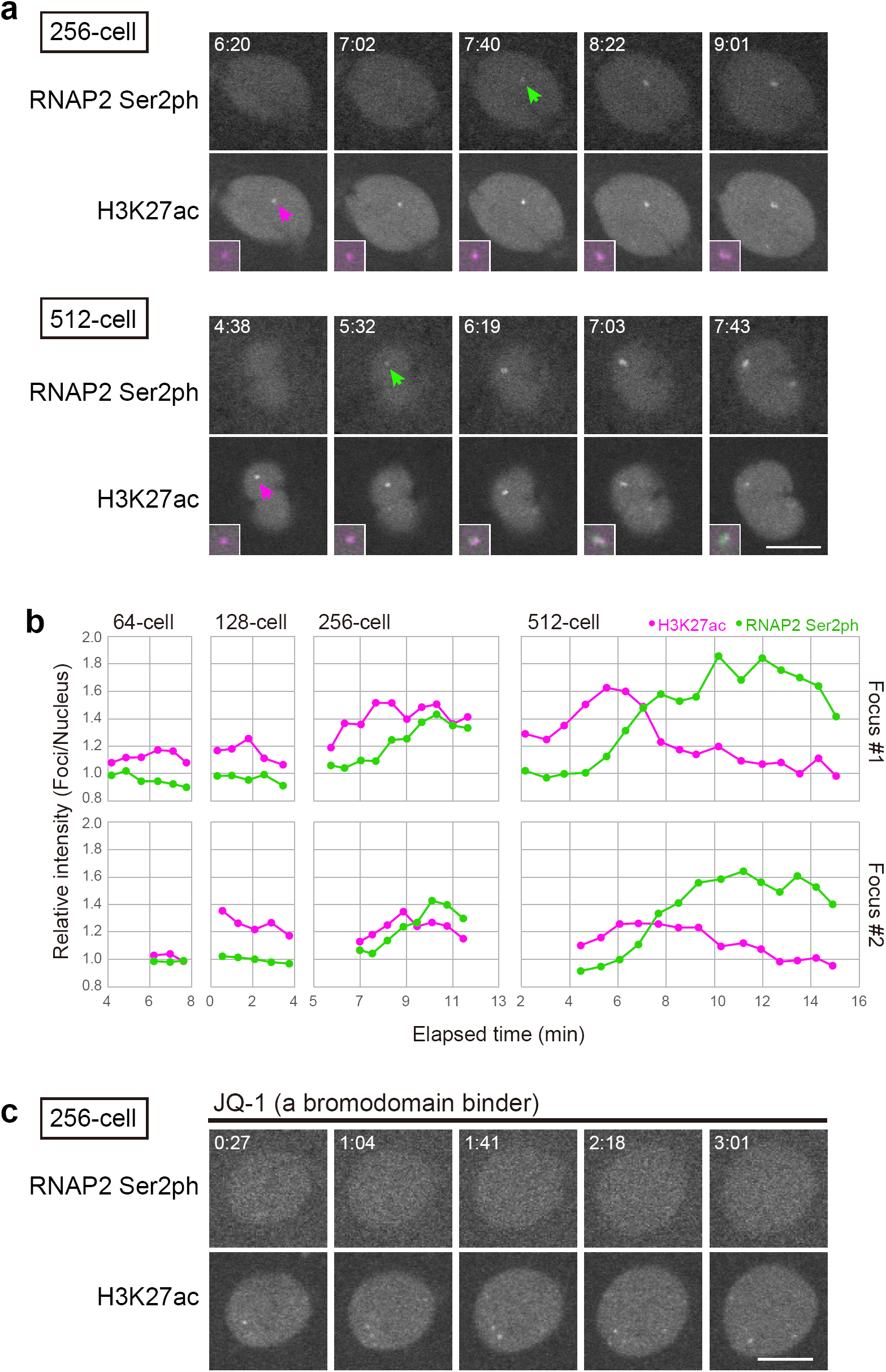
Dynamics of transcription foci at early stages. (a) Simultaneous visualization of RNAP2 Ser2ph and H3K27ac. RNAP2 Ser2ph accumulated at around H3K27ac foci. Magnified and merged images of foci are shown in insets (H3K27ac, magenta; RNAP2 Ser2ph, green). Elapsed time (m:ss) is indicated. (b) Relative intensity (Foci/Nucleus). (c) RNAP2 Ser2ph accumulation was inhibited by a BET domain binder, JQ-1. The injected embryo was soaked into 10 μM JQ-1 and imaged. Scale bars, 10 μm.

To investigate the relationship between H3K27ac and transcription, we used chemical compounds to inhibit RNAP2 transcription and acetyl reader proteins. The effect of RNAP2 inhibition by α-amanitin on the H3K27ac levels and its focus formation was negligible (Supplementary Fig. 7), indicating that active transcription is not required for H3K27ac accumulation. To investigate the effect of histone acetylation on transcription, we initially attempted to inhibit p300 histone acetyltransferase using its selective inhibitor C646 (Bowers et al., 2010). Nevertheless, soaking in 10 μM C646 did not affect H3K27ac level in embryos up to 3 hr, which may be due to poor permeability. We then used bromodomain and extra-terminal motif (BET) inhibitor, JQ-1. JQ-1 treatment deters BET family proteins from binding to acetylated histones and may inhibit transcription activation. In the presence of 10 μM JQ-1, RNAP2 Ser2ph was no longer concentrated in foci and H3K27ac foci appeared to become smaller and scattered (Fig. 3c). These results suggest that the binding of BET family proteins to H3K27ac is required for transcription activation and for enhancing H3K27 acetylation in the specific loci through a positive feedback. The pivotal role of p300 and BET family proteins in ZGA has also been demonstrated with the natural cell cycle or developmental stages. In this study, we demonstrated by Fab-based imaging that H3K27ac accumulation precedes RNAP2 activation during ZGA in living zebrafish embryos. Furthermore, even in the single nucleus, H3K27ac was concentrated prior to RNAP2 activation on miR-430 loci (Fig. 3b). These observations, together with inhibitor assays, suggest that H3K27ac and its reader proteins stimulate RNAP2-mediated transcription during ZGA.

## Conclusions

The method presented here is thus powerful to track the dynamics of RNAP2 and histone modifications during the development of living organisms. A further investigation of H3K27ac and RNAP2 Ser2ph during later stages will be interesting to provide a link between chromatin modification and transcription with nuclear organization (Hilbert et al., 2018; Kaaij et al., 2018), as RNAP2 Ser2ph distribution changes from the accumulation at two miR-430 foci to more scattered patterns. We anticipate that the method presented here, which does not involve genetically recombinant materials, can be applied to a variety of developmental processes in any model and non-model organisms.

## Methods

### Fabs

Fabs were prepared as previously described (Kimura and Yamagata, 2015) using mouse monoclonal antibodies specific to RNAP2 Ser2ph (CMA602/Pc26B5) (Stasevich et al., 2014), Ser5ph (CMA605/Pa57B7) (Stasevich et al., 2014), K7me2 (CMA612/19B4) (Dias et al., 2015), pan CTD (CMA601/C13B9) (Stasevich et al., 2014), H3K4me3 (CMA304/16H10) (Hayashi-Takanaka et al., 2011), H3K9ac (CMA310/19E5) (Hayashi-Takanaka et al., 2011), H3K27ac (CMA309/9E2) (Hayashi-Takanaka et al., 2011), H3K27me3 (CMA323/1E7) (Hayashi-Takanaka et al., 2011), H3K36me3 (CMA333/13C9) (Hayashi-Takanaka et al., 2011), H4K5ac (CMA405/4A7) (Hayashi-Takanaka et al., 2015), H4K16ac (CMA416/1B2) (Hayashi-Takanaka et al., 2015), H4K20me1 (CMA421/15F11) (Hayashi-Takanaka et al., 2015), and H4K20me2 (CMA422/2E2) (Hayashi-Takanaka et al., 2015). IgG was purified from hybridoma culture supernatant through a Protein A-Sepharose column (GE Healthcare), digested with agarose beads conjugated with Ficin (for IgG1) or Papain (for IgG2a and IgG2b) (Thermo Fisher), and applied to a Protein A-Sepharose column (GE Healthcare) to remove Fc and undigested IgG. The resulting Fabs were concentrated up to ∼1 mg/ml in PBS using a Microcon filter unit (Millipore) and labeled with NHS-ester of Alexa Fluor 488 (ThermoFisher), Cy3 (GE Healthcare), Cy5 (GE Healthcare), JF549 (Grimm et al., 2015), and JF646 (Grimm et al., 2015), as described previously (Kimura and Yamagata, 2015). Normal mouse IgG (Jackson ImmunoResearch) was also processed to yield dye-labeled control Fab.

### Morpholino for detecting miR-430

Morpholino antisense oligonucleotide (MO) to detect miR-430 was designed according to Hadzhiev et al. (2019) and 3’-primary amino-modified miR-430 MO (5’-TCTACCCCAACTTGATAGCACTTTC-3’) was obtained from Gene tools LLC and labeled with Cy3 NHS-ester (GE Healthcare). After the conjugation for 1 h at room temperature, Cy3-labeled miR-430 was separated from unconjugated dyes using a QIAquick Nucleotide Removal Kit (QIAGEN).

### Visualizing zebrafish embryogenesis

Zebrafish handling was operated according to the institutional guidelines of Tokyo Institute of Technology and HHMI Janelia Research Campus. Zebrafish (*Danio rerio*, AB) eggs were obtained from pairwise mating. Labeled Fabs (50-300 pg in ∼0.5 nl) and miR-430 Morpholino (1.8 fmol in ∼0.5 nl) were injected into the yolk of 1-cell stage embryos.

For collecting images using simultaneous multi-view light-sheet microscopy (Tomer et al., 2012), injected embryos were dechorionated and embedded in 2-mm or 3-mm glass capillaries (Hilgenberg GmbH) filled with 0.9% low melting temperature agarose (Type VII, Sigma Aldrich) prepared in zebrafish system water. The fully gelled agarose cylinder was extruded from the capillary to expose the embryo to the detection system. The capillary was then mounted vertically in the recording chamber containing zebrafish system water, so that the agarose section containing the embryo was mechanically supported by the glass capillary below with the animal pole of the embryo facing the microscope’s detection arm. Bi-directional scanned laser light sheets at a 488, 561, and 647 nm wavelengths were used for excitation. Fluorescence was detected using 525/50 nm band-pass, 561 nm long-pass, and 647 nm long-pass detection filters (Semrock), respectively. Imaging was performed using Nikon 16x/0.8 NA (Fig. 1b) or Zeiss 20x/1.0 NA (Fig. 1c) water immersion objectives and images were acquired with Hamamatsu Orca Flash 4.0 v2 sCMOS cameras. Time-lapse imaging was performed at a time interval of 1 min using the AutoPilot framework for automatic adaptive light-sheet adjustment. Image stacks of 197 planes encompassing the entire volume of the embryo with an axial step size of 2.031 µm were acquired for each time point. The lateral pixel sizes in the image data were 0.406 µm (Fig. 1b) or 0.325 µm (Fig. 1c).

For collecting images using a confocal microscope (FV1000; Olympus), injected embryos were incubated at 28°C until 4-cell stage. The embryos were dechorionated and embedded in 0.5% agarose (Sigma Aldrich, A0701) in 0.03% sea salt with the animal pole down on a 35-mm glass bottom dish (MatTek), which was set on to a heated stage (Tokai Hit) at 28°C. Fluorescent Images were acquired using an FV1000 (Olympus) operated by the built-in software FLUOVIEW ver.4.2 with a UPLSAPO 30x silicone oil immersion lens (NA 1.05), using 512×512 pixels, scan speed 2.0 μs pixel dwell time, zoom 1.0, and 4 μm z-interval (20-25 sections), with 488 nm (75 μW at the specimen), 543 nm (40 μW), and 633 nm (55 μW) laser lines.

### Inhibitors

α-amanitin (∼0.5 ng in ∼0.5 pl water; Merck Millipore) was injected into the embryo after injection of fluorescent probes. C646 (Calbiochem) and JQ-1 (BPS Bioscience) were added into embedding agarose at 10 μM.

### Quantification of fluorescence signals

To obtain time courses of nuclear-over-cytoplasmic intensity ratios, an existent pipeline for time-resolved, single nucleus level analysis was modified and extended (Joseph et al., 2017). In brief, the nuclear signal of the Cy5-labeled Fabs specific for histone H3 Lys9 acetylation (H3K9ac), which is a broad euchromatin mark, was used to segment and track individual nuclei during interphase. A manual review step on the basis of a graphical user interface was added to correct tracking errors. Based on single nuclei segmentation masks, the intra-nuclear and the cytoplasmic intensity was quantified using two derivative segmentation masks. The intensity data of single nuclei were exported to tabulated files that can be accessed and further analyzed using other software tools, e.g. Microsoft Excel. All image analysis procedures were implemented in MatLab, using the BioFormats Open Microscopy Environment importer bfMatlab (Goldberg et al., 2005) and are available as open source code (https://github.com/lhilbert/NucCyto_Ratio_TimeLapse). As the size of raw data (several hundreds of Gigabytes) makes permanent hosting impractical, raw data are available on request. For Fig 3b, single Z-planes where the sectional area of H3K27ac focus became maximum were selected at each time point from time-lapse images. The regions of interest (ROI) of 1 μm diameter circle just covering H3K27ac focus on the planes were obtained using Fiji (http://fiji.sc/). All intensities in the ROI were summed up.

## Acknowledgments

We thank all the member of Kimura lab at Tokyo Institute of Technology for experimental support and discussion, Yutaka Kikuchi and Akihiko Muto for the instruction of injection and imaging zebrafish embryos, and Luke Lavis for JF dyes. This work was supported by JSPS KAKENHI Grant Numbers 15K07157 and 17KK0143 (to Y.S.), 17H01417 and 18H05527 (to H.K), and NIH grant GM127538 (to TL). The Advanced Imaging Center at HHMI Janelia is jointly funded by the Gordon and Betty Moore Foundation and Howard Hughes Medical Institute.

## Competing interests

Authors declare no competing interests.

## Author contributions

Y.S. and H.K. conceived and designed the project. Y.S. performed most experiments and analyzed data. L.H. wrote MatLab code under the supervision by V.Z. and N.V. Y.W., J.M.H., T.-L. C. and P.K. constructed and operated the SiMView system. H.O. contributed for SiMView imaging. T.L. contributed to Morpholino experiments and edited the manuscript. N.V. contributed to zebrafish embryo work. Y.S. and H.K. drafted manuscript under the all other authors’ input, particularly, L.H., V.Z, and N.V.

## Supplementary Information

### Movie captions

**Supplementary movie 1. Live-imaging of zygotic genome activation.**

The embryo injected RNAP2 Ser2ph- and H3K9ac-Fab were imaged every minute from the 8-cell stage using SiMView. The maximum intensity projections of 198 z-sections with 2 μm intervals.

**Supplementary movie 2. Live-imaging of the embryo injected with RNAP2 Ser2ph-Fab and miR-430 MO.**

The maximum intensity projections of 198 z-sections with 2 μm intervals.

**Supplementary Figure 1.**
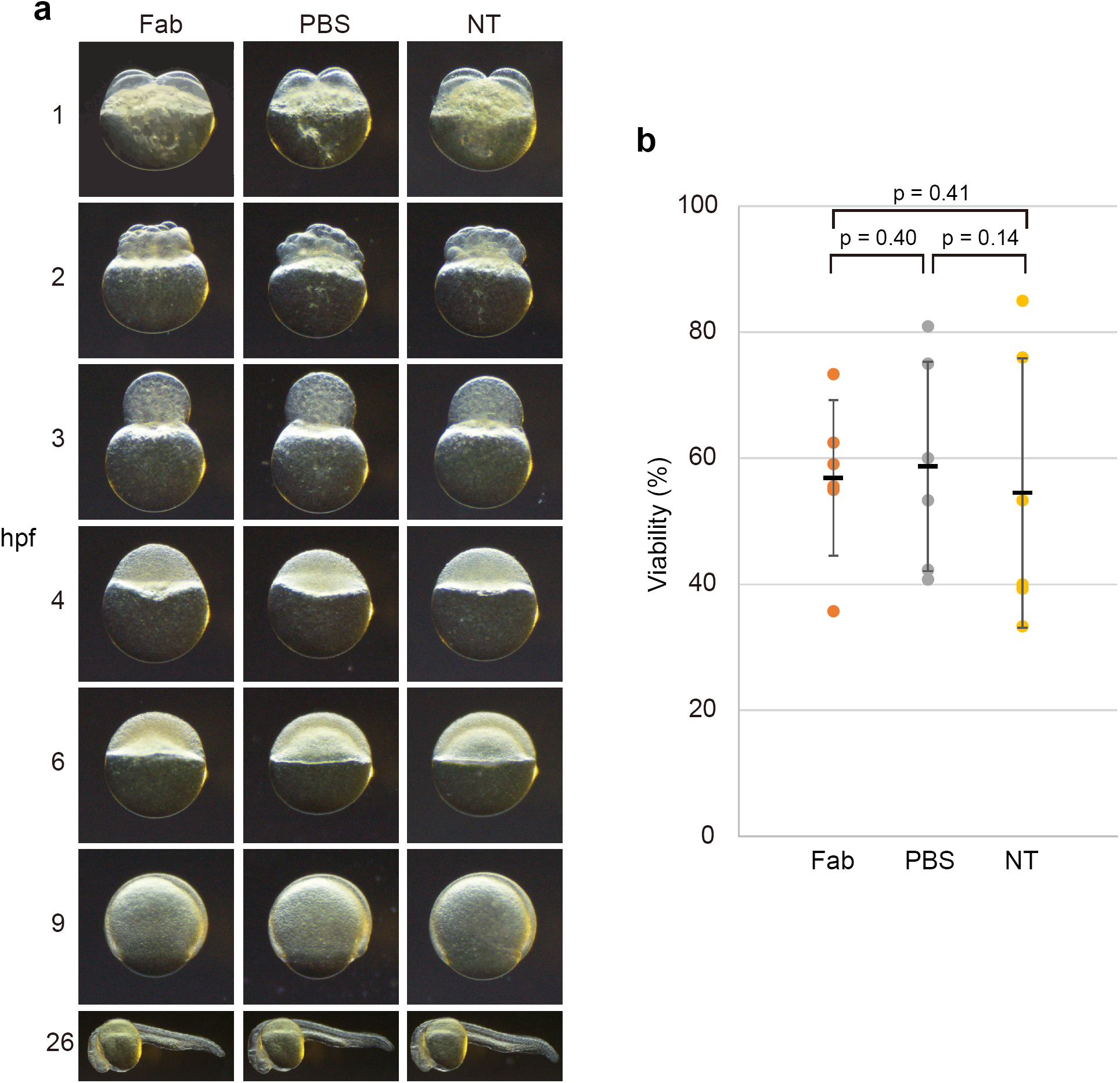
The influence of Fab injection on embryogenesis. (a) Images of the embryos injected with RNAP2 Ser2ph-Fab (Fab) or PBS are shown. (b) The percentages of viable embryos on the next day of injection from 5 independent experiments are plotted. NT, non-treated embryo.

**Supplementary Figure 2.**
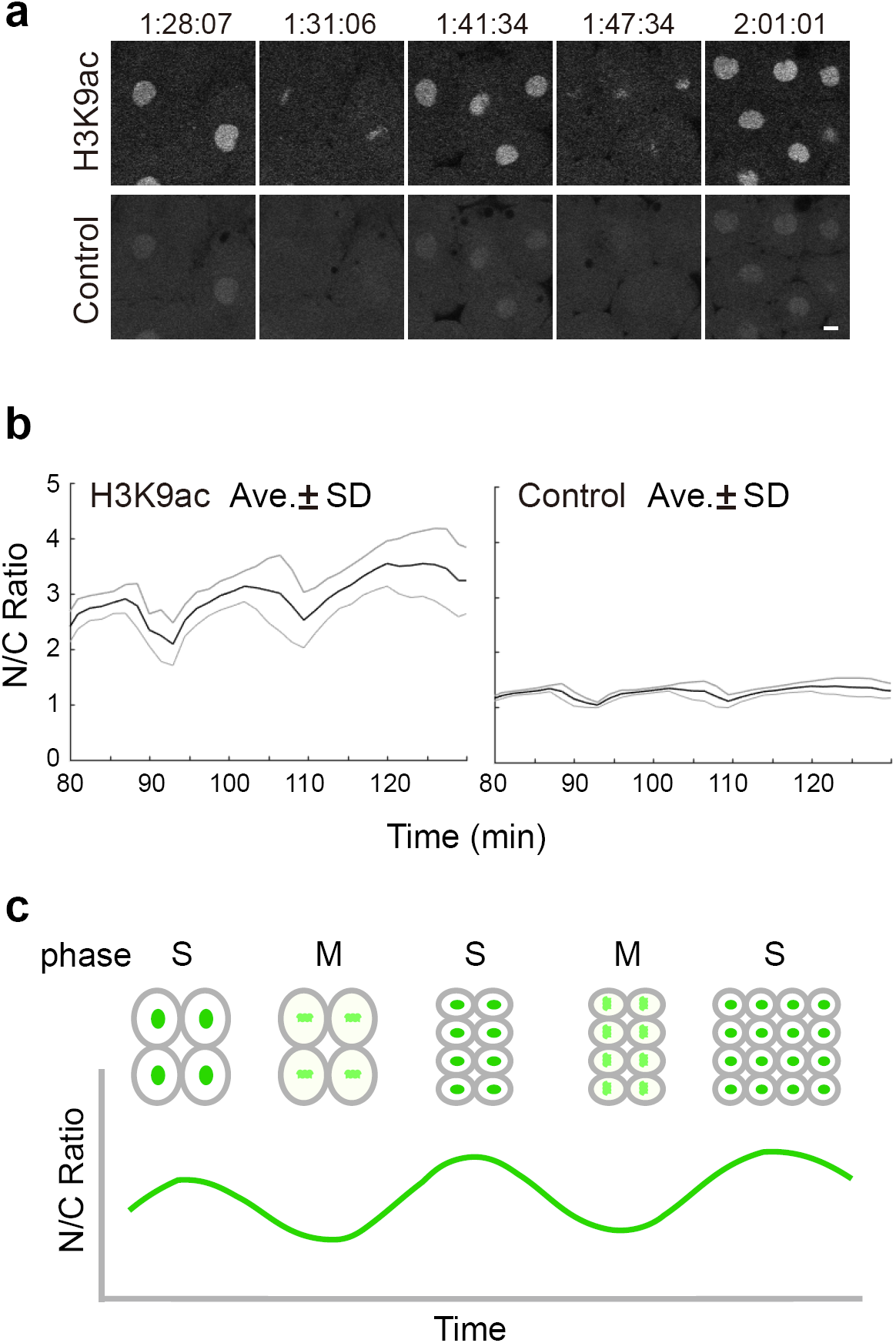
The intensity on mitotic chromosomes. (a) The embryo injected Fab specific to H3K9ac and non-specific Fab (Control) were imaged. Elapsed times (h:mm:ss) are indicated. Scale bar, 10 μm. (b) Nucleus/Cytoplasm intensity ratio of H3K9ac and Control Fab. Average ± Standard deviation is shown. (c) Schematic illustration for the oscillation of nuclear/cytoplasmic ratio during the cell cycle.

**Supplementary Figure 3.**
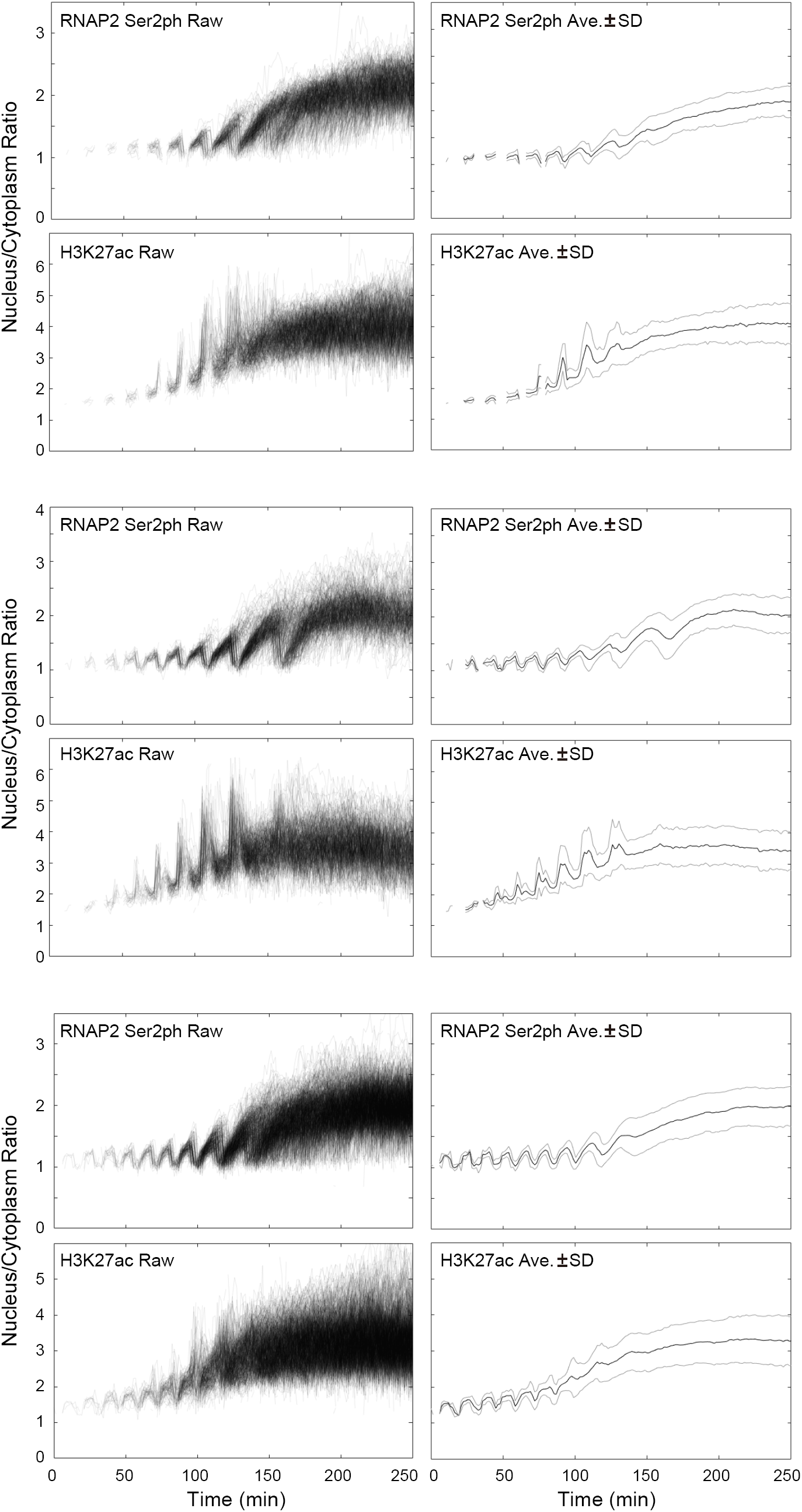
Nucleus/Cytoplasm intensity ratios of the embryos injected with RNAP2 Ser2ph- and H3K27ac-Fabs. Three different experiments are shown. A pair of graphs (e.g., the first and second, and the third and fourth) are data from a single embryo.

**Supplementary Figure 4.**
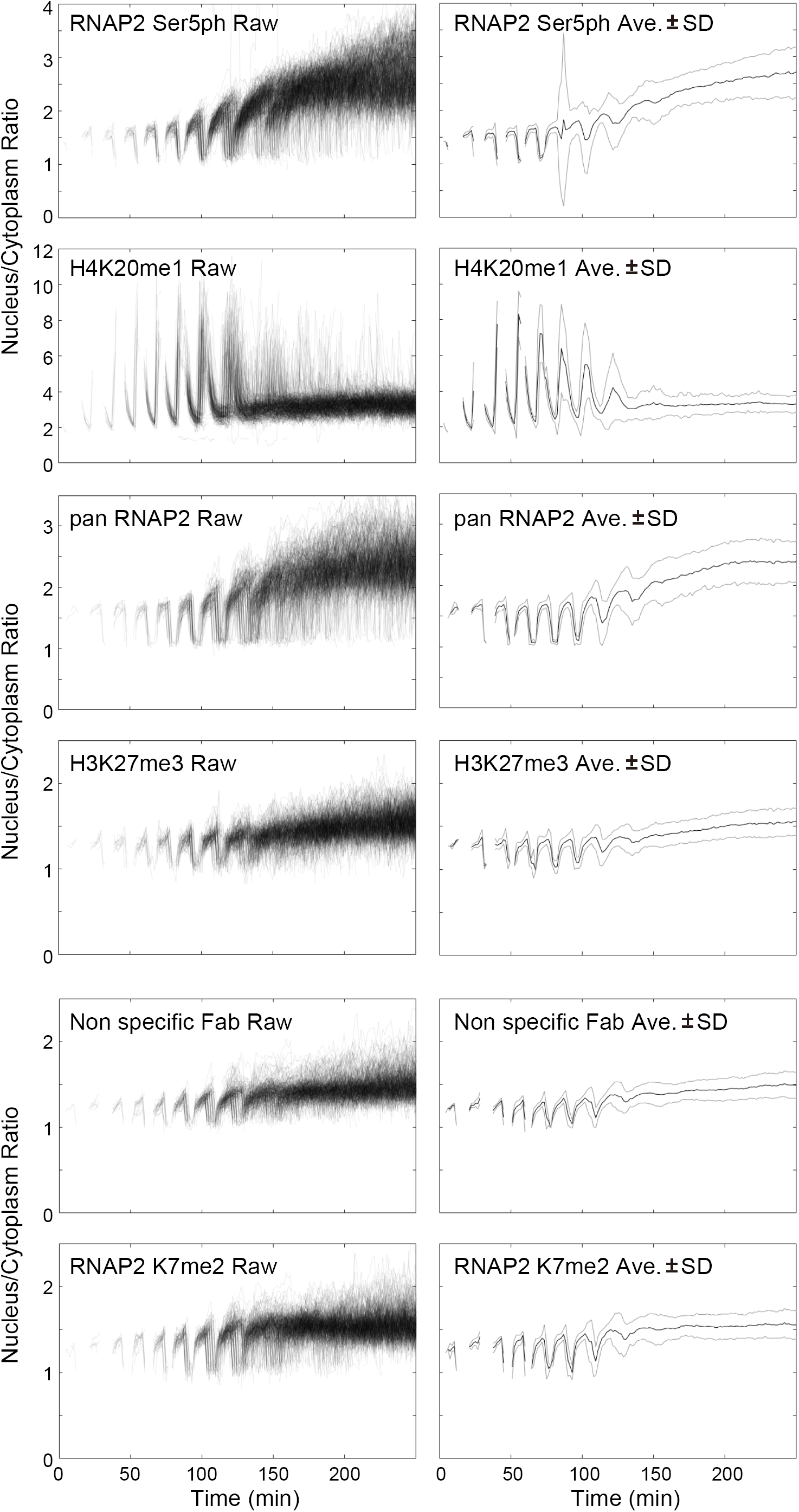
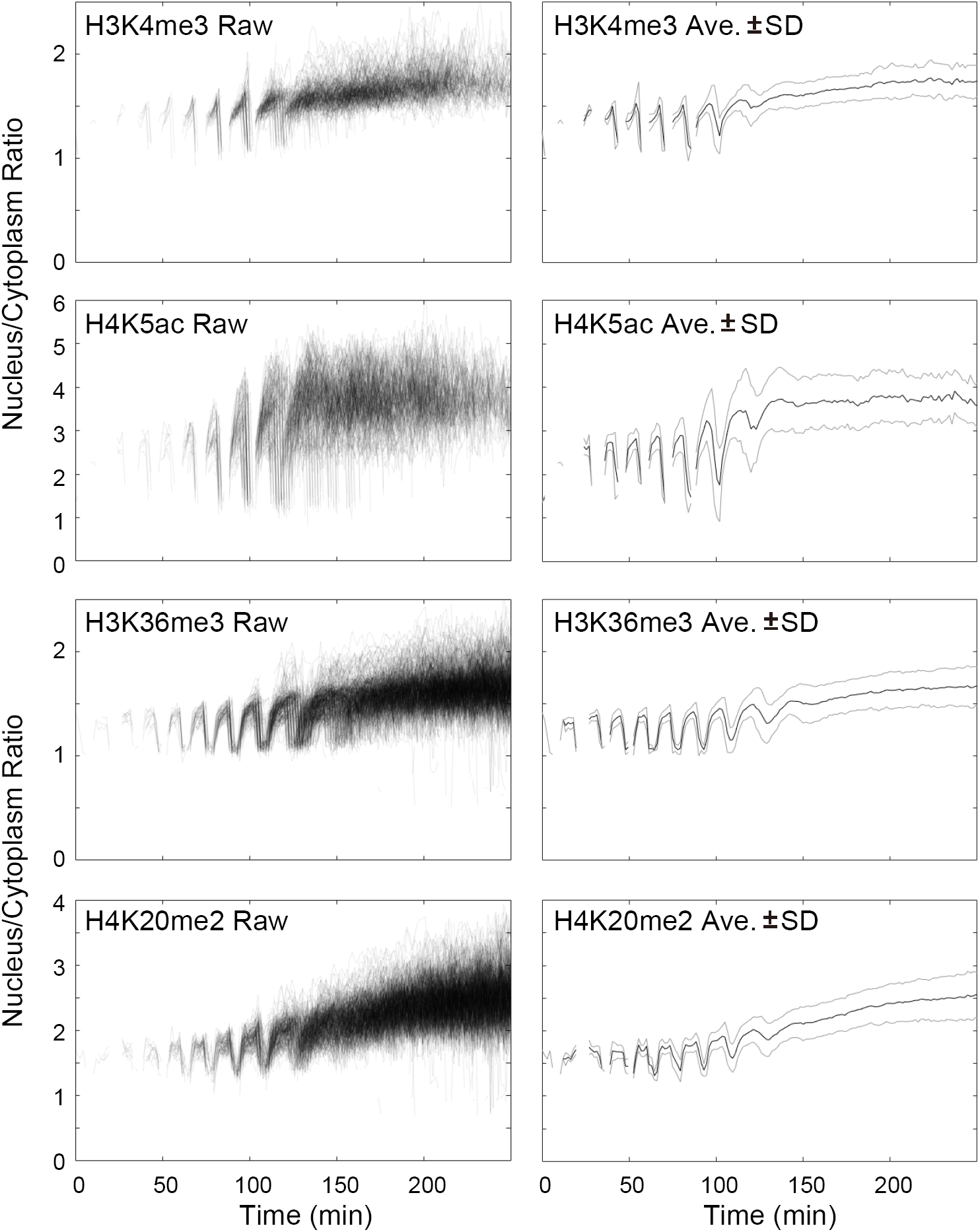
Nucleus/Cytoplasm intensity ratios of the embryos injected with Fabs specific to RNA polymerase II and histone modifications. A pair of graphs (e.g., the first and second, and the third and fourth) are data from a single embryo.

**Supplementary Figure 5.**
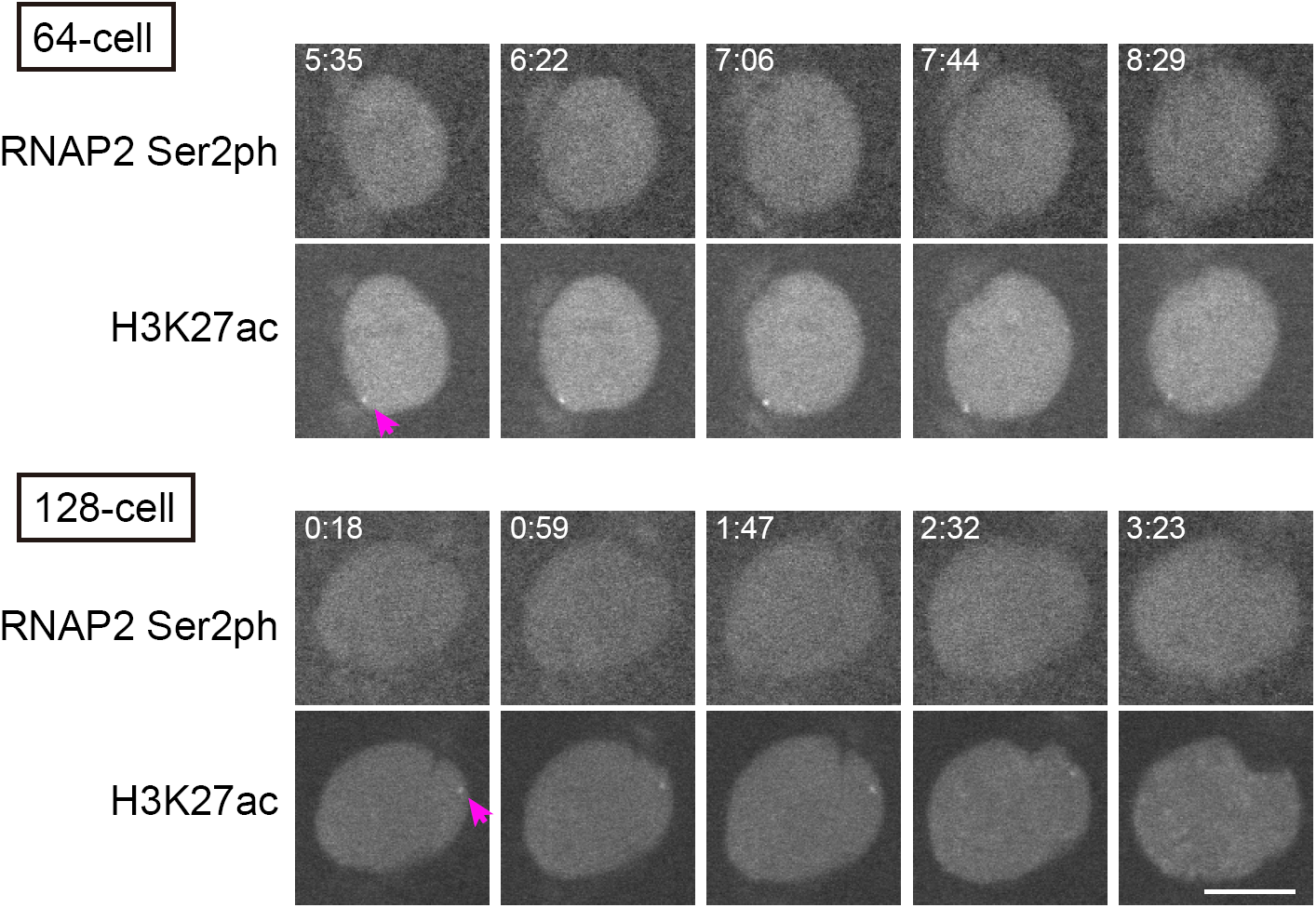
H3K27ac foci in early stage embryos. H3K27ac foci firstly appeared in interphase nuclei at the 64-cell stage, whereas RNAP2 Ser2ph accumulated in foci after the 256-cell stage. Elapsed times (m:ss) are indicated. Scale bar, 10 μm.

**Supplementary Figure 6.**
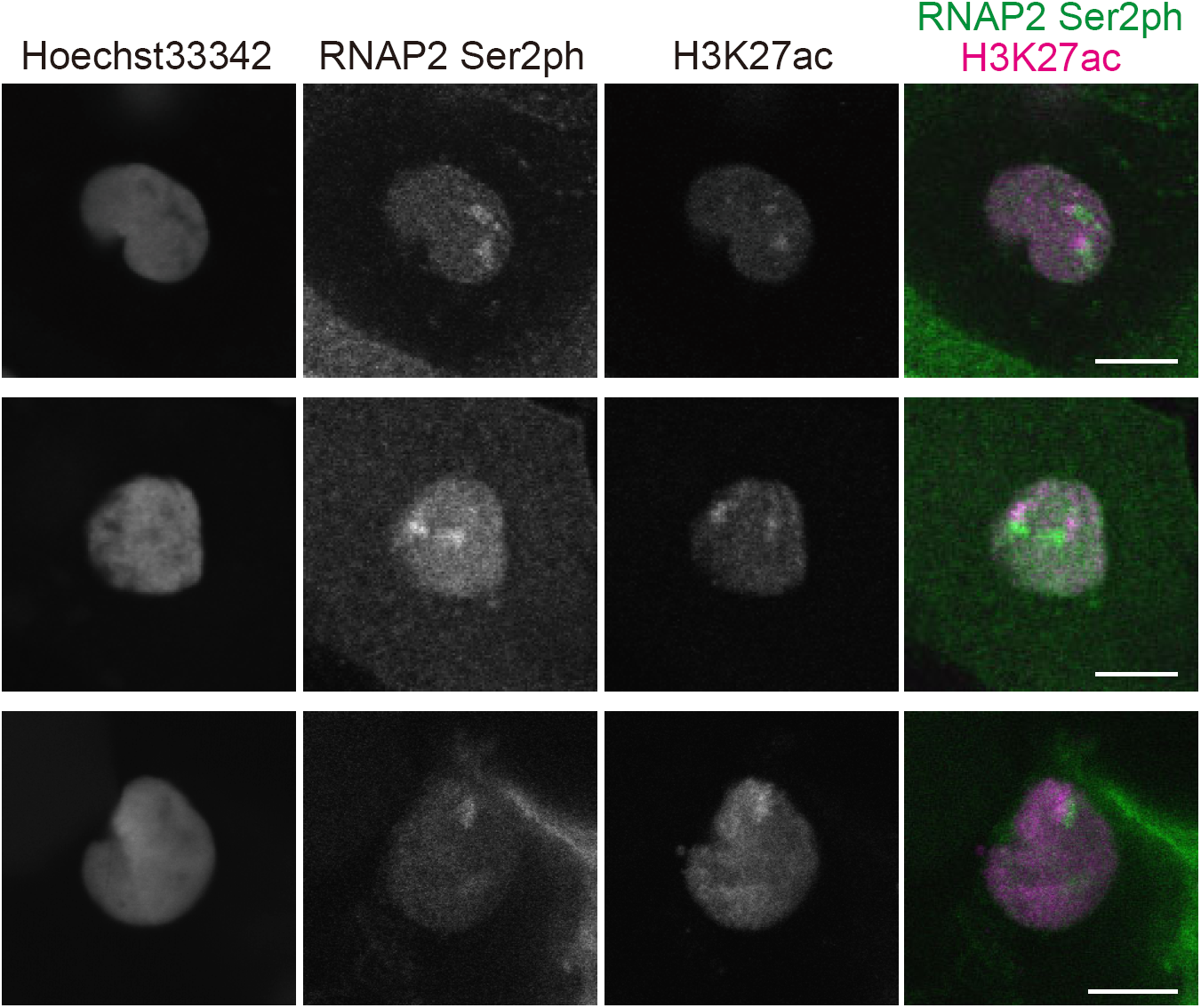
Co-localization of H3K27ac and RNAP2 Ser2ph. The embryos injected with RNAP2 Ser2ph-Fab were fixed at the 512-cell stage with fixation buffer (4% paraformaldehyde, 1% Triton X-100, 250 mM HEPES, 4°C overnight). After soaking in blocking One P (Nacalai Tesque) containing 0.1% Triton X-100 for 2 hour, embryos were stained with 10 μ g/ml anti-H3K27ac antibody (CMA309) and 1 μg/ml Hoechst33342. Images were taken with a confocal microscope (FV1000; Olympus). Scale bar, 10 μm.

**Supplementary Figure 7.**
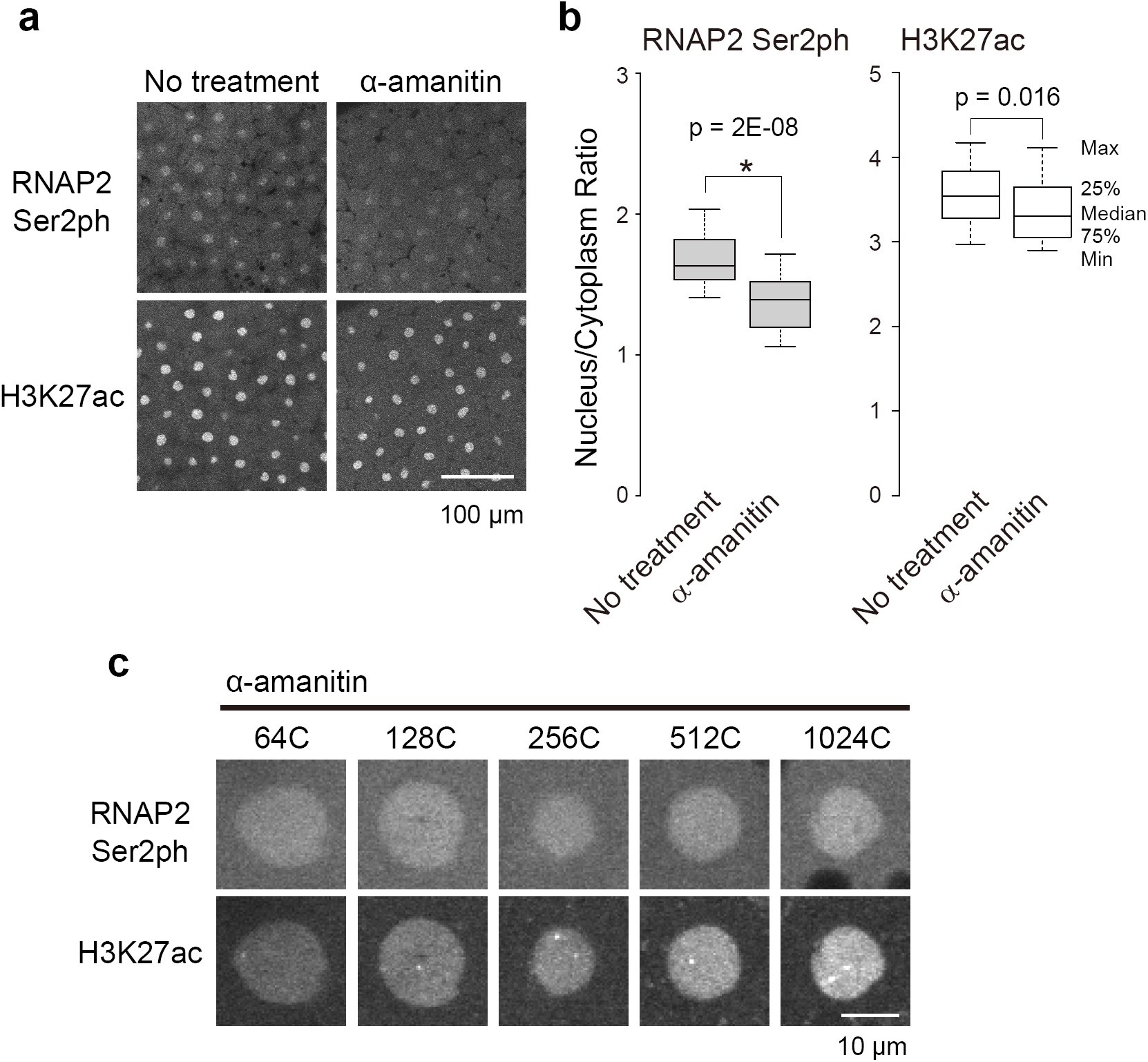
Inhibitors for RNA polymerase II activity did not affect H3K27 acetylation. (a) α-amanitin (∼0.5 ng) was injected into embryo. (b) Nuclear/Cytoplasm Ratios were measured (No treatment, n=32; α-amanitin, n=30). (c) The images of the 1k-cell stage embryo are shown. Scale bar, 100 μm.

